# Chromosome level reference genome for European flat oyster (*Ostrea edulis* L.)

**DOI:** 10.1101/2022.06.26.497633

**Authors:** Manu Kumar Gundappa, Carolina Peñaloza, Tim Regan, Isabelle Boutet, Arnaud Tanguy, Ross D. Houston, Tim P. Bean, Daniel J. Macqueen

## Abstract

The European flat oyster (*Ostrea edulis* L.) is a bivalve naturally distributed across Europe that was an integral part of human diets for centuries, until anthropogenic activities and disease outbreaks severely reduced wild populations. Despite a growing interest in genetic applications to support population management and aquaculture, a reference genome for this species is lacking to date. Here we report a chromosome-level assembly and annotation for the European Flat oyster genome, generated using Oxford Nanopore, Illumina, Dovetail OmniC™ proximity ligation and RNA sequencing. A contig assembly (N50: 2.38Mb) was scaffolded into the expected karyotype of 10 pseudo-chromosomes. The final assembly is 935.13 Mb, with a scaffold-N50 of 95.56 Mb, with a predicted repeat landscape dominated by unclassified elements specific to *O. edulis*. The assembly was verified for accuracy and completeness using multiple approaches, including a novel linkage map built with ddRAD-Seq technology, comprising 4,016 SNPs from four full-sib families (8 parents and 163 F1 offspring). Annotation of the genome integrating multi-tissue transcriptome data, comparative protein evidence and *ab-initio* gene prediction identified 35,699 protein-coding genes. Chromosome level synteny was demonstrated against multiple high-quality bivalve genome assemblies, including an *O. edulis* genome generated independently for a French *O. edulis* individual. Comparative genomics was used to characterize gene family expansions during *Ostrea* evolution that potentially facilitated adaptation. This new reference genome for European flat oyster will enable high-resolution genomics in support of conservation and aquaculture initiatives, and improves our understanding of bivalve genome evolution.

## Introduction

The European flat oyster *Ostrea edulis* (Linnaeus, 1758) is a bivalve mollusc within Ostreidae (‘true oysters’). This species is a native of Europe, naturally distributed from 65 degrees North in Norway to 30 degrees North in Morocco, along the North-Eastern Atlantic, and also the entire Mediterranean basin (Thorngren et al., 2019). Introductions of *O. edulis* in the 19^th^ and 20^th^ centuries for aquaculture resulted in the establishment of natural beds in many regions across the world, including North America, New Zealand, Australia, and Japan (Bromley et al., 2016). *O. edulis* can reach sizes exceeding 20cm and has a life span up to 20 years (Bayne, 2017). This species is a protandrous hermaphrodite that can change sex within a spawning season, and unlike the more widely cultured Pacific oyster *Crassostrea gigas*, brood their larvae in the inhalant chamber for several days before release (Suquet et al., 2018). *O. edulis* exhibits extensive physiological plasticity across its range, for example the temperature at which spawning occurs (11-25°C degrees) and the duration of the spawning period (from 1-2 months, to year round) (Bromley, 2015; Bromley et al., 2016).

*O. edulis* has been an integral part of human diets in Europe for centuries, with evidence for its collection and consumption since at least Roman times. Furthermore, it is thought >700 million oysters were consumed in London alone during 1864 (Pogoda, 2019a). However, overfishing and anthropogenic activities have driven a collapse of *O. edulis* stocks throughout its natural range (Pogoda, 2019b; Merk et al., 2020). The past 40 years has witnessed a further decline in production, with a peak of 32,995 tonnes in 1961 dropping by >90% to 3,120 tonnes by 2016 (FAO, 2020). Human impacts are widely cited as the primary reason for this decline, including habitat destruction, overexploitation, the introduction of non-native species competing for *O. edulis* habitats (Grizel & Héral, 1991; Vera et al., 2019), and the emergence/spread of diseases associated with translocations (Bromley et al., 2016). Key parasites associated with flat oyster population declines include the protist *Marteilia refringens* and the haplosporidian protozoan parasite *Bonamia ostreae*, which causes bonamiosis, for which no effective control methods exist (Sas et al., 2020). Large scale restoration efforts exemplified by the Native Oyster Restoration Alliance (NORA; https://noraeurope.eu/) are targeting re-stocking of *O. edulis* at high densities and developing sustainable populations. However, these efforts are strongly hampered by parasitic disease, especially bonamiosis (Engelsma et al., 2010; Pogoda et al., 2019a). While using animals from *Bonamia* free regions offers a potential short-term solution for restoration and aquaculture efforts, understanding the genetic basis for natural parasite resistance (Sas et al., 2020) will enable selective breeding to enhance *Bonamia* resistance and permanently reduce disease incidence in farmed and wild populations.

Several studies have applied genetic and genomic tools to study *O. edulis* in the absence of a reference genome assembly. Such work has been strongly targeted towards understanding bonamiosis, either by identifying candidate quantitative trait loci (QTL) and genetic outliers linked to *Bonamia* resistance (Lallias et al., 2009; Harrang et al., 2015; Vera et al., 2019) or by studying gene expression responses to *Bonamia* infection (Pardo et al., 2016; Ronza et al., 2018). SNP genotyping arrays with low (Lapègue et al., 2014) and medium (Gutierrez et al., 2017) density have also been developed for genetics applications. The lack of a high-quality reference genome in *O. edulis* however, contrasts with the situation in the commercially valuable Pacific oyster *C. gigas* (Peñaloza et al., 2021; Qi et al., 2021) and is a current limitation for the research community. An annotated genome for *O. edulis* will enable genetics research in many directions supporting conservation and aquaculture, revealing the physical location of genetic variation with respect to genes and genomic features, and offering an essential foundation for functional genomics. A reference genome will also support our understanding of *O. edulis* evolution and environmental adaptation, through comparisons with other bivalve species.

Bivalve genome assembly has classically been hampered by genetic complexities including high heterozygosity and repeat content (Davison & Neiman, 2021), along with the challenge of extracting pure high-molecular weight DNA (Adema, 2021). However, recent advances in long-read sequencing technologies have enabled high quality genome sequences to be generated for multiple bivalves, including *C. gigas* (Peñaloza et al, 2021; Qi et al, 2021), the scallop *Pecten maximus* (Kenny et al., 2020) and hard clam *Mercenaria mercenaria* (Song et al., 2021; Farhat et al., 2022). Here, we integrated multiple sequencing technologies to assemble and annotate a highly contiguous chromosome-level genome assembly for an *O. edulis* individual from the UK, which was confirmed for accuracy by comparison to a novel linkage map for *O. edulis*, and high-quality genome assemblies for several bivalve species. Comparative genomics inclusive of diverse bivalve species allowed us to define gene copy expansions in the *Ostrea* lineage. The high-quality reference genome reported here, and an independent *O. edulis* assembly reported for an individual from a distinct European population in the same issue of this journal by Boutet et al. (2022), will support ongoing conservation and aquaculture initiatives for the European flat oyster, while improving our comparative understanding of genome evolution and adaptation in the *Ostrea* lineage.

## Materials and Methods

### Data availability

The genome assembly generated in this study along with all raw sequencing data used in assembly and annotation (Oxford Nanopore reads used for contig assembly, Illumina paired-end reads used for contig/scaffold polishing, Dovetail® Omni-C™ paired end reads used for contig scaffolding, RNA-Seq paired-end reads from 8 tissues used for genome annotation) is available through NCBI under the Bioproject PRJNA772111. The genome annotation and large Supplementary Tables that are not available within the Supplementary Information are available through Figshare (https://doi.org/10.6084/m9.figshare.20050940).

### Sampling and sequencing

A single unsexed adult *O. edulis* individual sourced from Whitstable (England, UK) through a commercial supplier (Simply Oysters) was used for all DNA and RNA sequencing performed in this study, as described below. The oyster was depurated in clean seawater for at least 42 hours before sampling. Samples of gill, mantle, heart, white muscle, striated muscle, digestive gland, labial palp, and gonad were flash frozen using liquid nitrogen and stored at -80°C. High molecular weight DNA was extracted from gill using a cetyltrimethylammonium bromide (CTAB) based extraction method and used to generate short and long-read sequencing libraries. DNA purity was confirmed using a Nanodrop 1000 (Thermo Fisher Scientific). DNA integrity was initially assessed using a Tapestation 4200 (Agilent Technologies). The DNA was purified using Ampure beads (Beckman Coulter™), sheared to a length of ∼35 Kb using a Megaruptor® (Diagenode) and size selected in the 7-50 Kb range on a Bluepippin system (Sage Science) with a 0.75% cassette. The resulting DNA was sequenced on four PromethION flow cells (FLO-PRO002), with basecalling performed using Guppy version 3.2.6+afc8e14. Short-read libraries with an insert size of 350 bp were generated using the same DNA with an Illumina TruSeq DNA library kit, prior to sequencing on an Illumina NovaSeq 6000 by Novogene Ltd (UK) with a paired-end 150 bp configuration. An Omni-C™ library was generated from gill tissue by Dovetail Genomics (Santa Cruz, USA) and sequenced on an Illumina HiSeq X with a paired-end 150 bp configuration.

For RNA-Seq library generation, total RNA was extracted for the eight tissues using a Trizol based method, before DNAase treatment. RNA integrity was assessed using agarose gel electrophoresis and Bioanalyszer 2100 (Agilent). RNA purity was confirmed via a Nanodrop 1000 system. Illumina TruSeq mRNA libraries were prepared for each sample and sequenced on an Illumina NovaSeq 6000 with a paired-end 150 bp configuration by Novogene Ltd (UK).

### Genome assembly and scaffolding

Genome size and heterozygosity were estimated using a k-mer approach. The Illumina data was quality assessed using FastQC v0.11.8 (Andrews, 2010), trimmed using TrimGalore 0.4.5 (Krueger, 2015) (quality score >30, minimum length > 40 bp) and processed through Meryl v1.3 (Rhie et al., 2020) to generate a k-mer count database (k = 20), which was used to generate a k-mer histogram. The histogram data was used as an input to Genomescope 2.0 (Ranallo-Benavidez et al., 2020) to estimate genome size and heterozygosity.

Contig assembly was performed using the nanopore data with the repeat graph based assembler Flye 2.7-b1585 (Kolmogorov et al., 2019). Three contig assemblies were generated (*OE_F1, OE_F2, OE_F3*) setting the –*minimum-overlap* parameter to ‘5,000’, ‘10,000’, and ‘auto’, respectively, with all other parameters default. In parallel, the raw nanopore reads were error corrected using the *correct* module within Necat v0.0.1 (Chen et al., 2021b). The corrected reads were also assembled to contigs using the overlap based assembler wtdbg2 2.5 (Ruan & Li, 2020) with default parameters, generating the assembly *OE_RB1*. The Flye and wtdbg2 assemblies were passed through pseudohaploid (https://github.com/schatzlab/pseudohaploid) to purge un-collapsed haplotigs. The three purged Flye assemblies (*OE_F1_purged, OE_F2_purged, OE_F3_purged*) were merged using Quickmerge v0.3 (Chakraborty et al., 2016) setting the parameters *-hco 5*.*0 -c 1*.*5 -l n -ml m* to generate a merged assembly (*Flye_Merged*). Finally, the *Flye_Merged* and haplotig purged wtdbg2 (*OE_RB1_purged*) assemblies were merged using Quickmerge v0.3 (as above) to generate a final contig assembly (*OE_contig_v1*), which was polished for two rounds using quality-trimmed Illumina data with Pilon v1.24 (Walker et al., 2014) (*OE_contig_pilon_v1*).

The polished contig assembly was scaffolded by Dovetail Genomics using HiRise (Putnam et al., 2016) with the Omni-C™ proximity ligation sequencing data used to orient and link the contigs using 3D contact information. The top 10 super scaffolds with the HiRise assembly were > 40Mb and matched the expected *O. edulis* karyotype (n=10) (Thiriot-Quiévreux, 1984; Leitao et al., 2002; Horváth et al., 2013) (Figure 1a). The next two largest scaffolds (scaffolds 11 and 12, respective sizes: 13.5 and 9.4 Mb) were not assigned to one of the 10 super scaffolds despite their large size, which led us to hypothesise these regions belonged to the 10 super-scaffolds, yet had not been scaffolded by HiRise. In support of this hypothesis, visualisation of the 3D contact information using Juicebox (Durand et al., 2016a) revealed 3D contacts between HiRise scaffold 11 and scaffold 6 and between HiRise scaffold 12 and scaffold 1 (Supplementary Figure 1). To confirm these interactions, we repeated contig scaffolding with the Omni-C™ data using Juicer (default parameters) (Durand et al., 2016b) and the resultant assembly was aligned and compared with the HiRise assembly using QUAST (Gurevich et al., 2013). Visualisation of QUAST alignments in Icarus (Mikheenko et al., 2016) confirmed the locations of scaffolds 11 and 12 within super-scaffolds 6 and 1, respectively (Supplementary Figure 1). Manual integration of these scaffolds in the HiRise assembly was performed using Scaffolder (Barton & Barton, 2012). Following this work, super-scaffold 6 became the second largest super-scaffold, and was therefore renamed to be super-scaffold 2, and this annotation is used hereafter. The resulting scaffolds were polished for one round using Pilon, leading to the final assembly used in all downstream work (*OE_Roslin_V1*).

**Figure 1.**
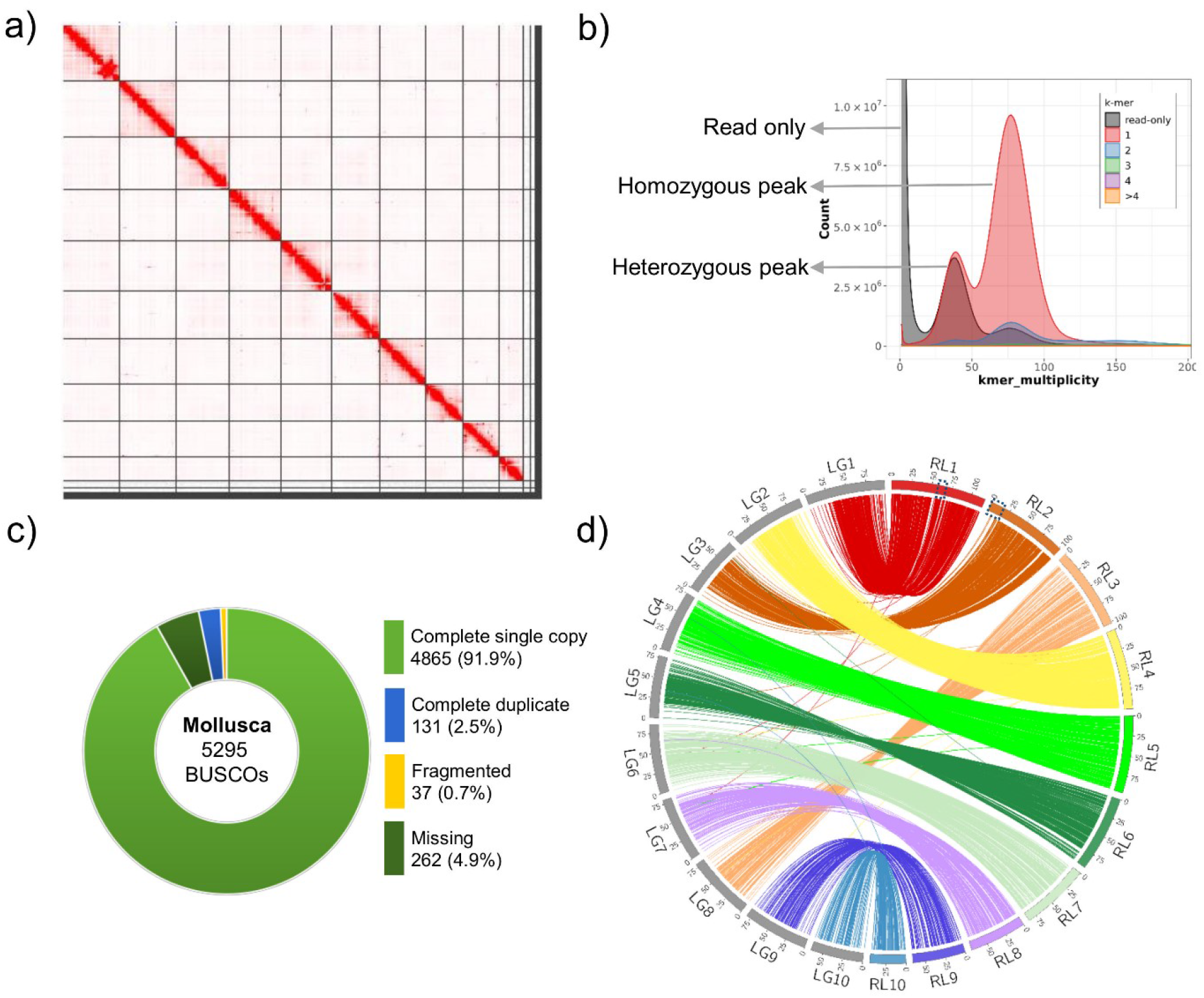
*OE_Roslin_V1* assembly quality evaluation. a) Omni-C contact map highlighting the top 10 super-scaffolds generated by HiRise. The contact map was visualised using Juicebox (Durand et al., 2016a). b) Merqury k-mer copy number spectrum plot for the curated genome assembly. Nearly half of the single copy k-mers (black region) were missing from the heterozygous peak, indicating efficient purging of haplotigs from the final assembly. k-mers missing from the assembly (black region in the homozygous peak) indicates bases present in the Illumina data missing from the assembly. c) BUSCO scores for the final scaffolded *OE_Roslin_V1* assembly (mollusca_odb10 database). d) Circos map highlighting the concordance between the 10 super-scaffolds (RL1 to RL10) and linkage groups (LG1 to LG10). Blue dotted squares within super-scaffolds 1 and 2 highlight the manual scaffolding performed on the basis of 3D contact information in the Omni-C data (Supplementary Figure 1).

### Genome quality evaluation

*OE_Roslin_V1* was screened for the presence of DNA contamination from other taxa using Blobtools v1.1.1 (Laetsch & Blaxter, 2017b) and for misassembly errors using Inspector v1.0.2 (Chen et al., 2021c). Structural errors identified in the genome were corrected using the Inspector-correct.py step. The raw nanopore reads were mapped back to the *OE_Roslin_V1* assembly using minimap2 (Li, 2018) (parameter *-ax map-ont*) to check for assembly completeness. The genome assembly was compared to a novel linkage map to confirm the accuracy of scaffolding using the chromatin proximity Omni-C™ data (see later section). Assembly quality and efficiency of haplotig purging was evaluated by generating a copy number spectrum plot (tracking the multiplicity of each k-mer in the read set, revealing the number of times it is found in the genome assembly) using Merqury v1.3 (Rhie et al., 2020). Gene completeness was evaluated against a set of 5,295 benchmark molluscan orthologous genes (*mollusca_odb10*) using BUSCO v4.1.4 (Simão et al., 2015). We mapped paired end Ilumina data from the same individual to the finished genome assembly using the minimap2 (Li, 2018) (parameter –ax sr). SAMtools (Danecek et al., 2021) was used to extract mean mapping depth values across the entire genome at 100kb intervals. GC content across the genome was retrieved using BEDTools v2.29.2 (Quinlan & Hall, 2010) at an interval of 500kb. The mean mapping depth and GC content data was plotted as a circos plot using the package Circlize 0.4.14 (Gu et al., 2014).

### Genome annotation

*De novo* repeat prediction was carried out using RepeatModeler v2.0.2 (Flynn et al., 2020). RepeatMasker v4.1.1 (Smit et al., 2015) was used for repeat masking with two databases: i) RepBase-20170127 (Jurka et al., 2005) for Pacific oyster (set using parameters “*-s Crassostrea gigas* –e ncbi”) and ii) the *de novo* repeat database generated by RepeatModeler. Gene model prediction was carried out on the repeat masked assembly using Funannotate v1.8.7 (Palmer, 2017) after using the Funannotate clean module. Following this, the RNA-seq reads were aligned to the genome using minimap2 v2.21-r1071 (Li, 2018). Proteins sequences for *C. gigas* and *C. virginica* from the UniProt database were aligned using Diamond v2.0.9 (Buchfink et al., 2021) and the resultant BAM files utilized for gene model prediction. PASA v2.4.1 (Haas et al., 2003) was then used to predict an initial set of high-quality gene models, which were used to train and run Augustus v3.3.32 (Stanke et al., 2006), SNAP (Korf, 2004) and GlimmerHMM v3.0.4 (Majoros et al., 2004). 40,283 high quality gene models were automatically extracted from the *ab-initio* predictions before passing all the data to EVidenceModeler v1.1.1 (Haas et al., 2008) for a final round of gene model prediction. Gene models <50 aa in length (n=2), spanning gaps (n=2), and transposable elements (n=5,330) were filtered by Funannotate before the retained gene models underwent UTR prediction using PASA. Functional annotation was performed using the annotate step within Funannotate. Interproscan (Jones et al., 2014) was used to annotate predicted gene products against the following databases: Pfam (El-Gebali et al., 2019), Panther (Mi et al., 2021), PRINTS (Attwood et al., 2012), Superfamily (Pandurangan et al., 2019), Tigrfam (Haft et al., 2013), PrositeProfiles (Sigrist et al., 2013), and Gene Ontology (GO) (The Gene Ontology Consortium, 2019). eggNOG-mapper v2.1.2 (Huerta-Cepas et al., 2017) was used to add functional annotation using the fast orthology assignment algorithm. BEDTools v2.29.2 (Quinlan & Hall, 2010) was used to extract data on genic content, gene density, classified repeats across unclassified repeats across the entire genome at a regular interval of 500kb, all this data was incorporated into a circos plot using the package Circlize 0.4.14 (Gu et al., 2014).

### Additional validation of manually incorporated scaffolds

As mentioned above, two scaffolds were manually incorporated into the HiRise assembly (also see Results). To confirm the validity of these scaffolds beyond the quality assessments described above, we confirmed the genes present in these regions were: i) of oyster origin, and ii) showed bioactivity comparable to other regions along the same chromosomes. Firstly, we retrieved the coding sequence of all genes predicted within the manually-incorporated and remaining regions of super-scaffolds 1 and 2, which were subjected to BLASTn (Altschul et al., 1997) searches the Pacific oyster genome (NCBI accession: GCA_902806645.1) and an independent Flat oyster genome (Boutet et al, 2022). The BLASTn cut-off was <1e-20 with remaining parameters default. Secondly, RNA-Seq data from heart, striated muscle and gonad were mapped to the genome assembly using STAR (Dobin et al., 2013) with default parameters. Mean RNA-Seq mapping depth for all gene models along super-scaffolds 1 and 2 was retrieved using SAMtools. Graphs comparing statistics between the manually-incorporated and remaining regions of super-scaffolds 1 and 2 were generated using ggplot2 (Wickham, 2016).

### Linkage map construction

Four oyster full-sibling families (n=171 individuals representing 8 parents and 163 F1 offspring) were used to build a novel linkage map for *O. edulis*. The families were produced in the Porscave hatchery (Lampaul-Plouarzel, Brittany, France). DNA was extracted from the parents and the offspring using a standard phenol-chloroform-isoamyl alcohol (PCI; 25:24:1, v/v) protocol. After two washes with PCI, DNA was precipitated overnight with absolute ethanol at -20°C, centrifuged, washed with 70% ethanol, dried and suspended in PCR-grade water. All DNA samples were run in a 1% agarose 1X TBE gel and quantified using a Qubit fluorometer (Thermo Fisher Scientific) with a high-sensitivity dsDNA quantification kit (Invitrogen) according to the manufacturer’s instructions. Double-digest RAD-seq (ddRADSeq) libraries were produced for every sample following Brelsford et al. (2016). Briefly, for each individual, 200 ng of genomic DNA was digested using four different enzyme combinations (KasI/AciI, KasI/HpyCH4IV, KasI/MspI and PstI/MseI) (New England Biolabs). Barcoded adaptors were ligated to the digested DNA fragments and purified using Nucleo Mag NGS Clean-up and Size Select Kit (Macherel-Nagel). 8µl of purified template was used for enrichment and Illumina indexing by PCR using Q5 hot start DNA polymerase (New England Biolabs) (PCR conditions: 98°C 30s, 15 cycles 98°C 10s, 60°C 20s, 72°C 30s). A final elongation was done by adding buffer, dNTPs and primers for 15 min at 72°C. PCR products were run in a 1% agarose 1X TBE gel, quantified using a Qubit fluorometer with a high sensitivity dsDNA quantification kit (Invitrogen) and then pooled in equal proportions into two separate libraries. A 300-800 bp size selection was performed using a 1.5% agarose cassette in a Pippin Prep instrument (Sage Science). Each fraction was run through a DNA chip in a Bioanalyser (Agilent) to determine mean fragment size. The libraries were pooled at equimolar concentration and sequenced on one lane of a NovaSeq 6000 by Novogene Ltd (USA).

Raw reads were cleaned and demultiplexed with Stacks v2.5.4 (Catchen et al., 2013; Rochette et al., 2019). To avoid reference bias in the quality assessment of the genome assembly, SNP discovery and genotyping was performed using a *de novo* approach. To identify optimal parameter settings, two Stacks parameters were evaluated: (M) the maximum number of nucleotide mismatches allowed between stacks (or putative alleles) and (m) the minimum number of identical reads used to form a stack. For a subset of 12 samples, values of M were varied from 2-9, while parameter m was fixed to either 3 or 5. The final optimal parameter settings (m = 3, M = 4) were chosen as the combination of values that resulted in the highest number of polymorphic loci shared across 80% of the individuals (r80 rule) (Paris et al., 2017). Variants were called from the *de novo* assembled data if the locus was present in more than 80% of the individuals (-r 0.8), after removing sites with an observed heterozygosity higher than 0.7 (--max_obs_het 0.7). Genotyping in Stacks resulted in a total of 28,447 assembled loci, with an average depth across polymorphic sites of 79x and 29x in the parental and offspring samples, respectively. The consensus sequences of the catalogued loci were exported and the first 150bp mapped to *OE_Roslin_V1* using BWA v0.7.8 (Li & Durbin, 2009). Variants within ddRAD loci with a mapping quality (MAPQ) >4 were retained for subsequent analysis. Among these loci, 98% (24,079 out of 24,522) were uniquely mapped to the *O. edulis* genome and had the same or fewer mismatches than the default value (MAPQ ≥ 25) (Menzel et al., 2013).

Further quality control (QC) filters were applied to the genotype data in Plink v1.9 (Chang et al., 2015). Markers and individuals with excess missing data (>10%) were discarded. A principal component analysis revealed that seven individuals separated from their family cluster (Supplementary Figure 2). Upon closer inspection, their high levels of Mendelian errors (>100 errors) suggested they had been mislabelled and were therefore removed from the dataset. After QC-filtering, 15,373 SNPs genotyped across 8 parents and 163 offspring were available for the construction of a linkage map using Lep-Map2 and Lep-Map3 (Rastas et al., 2016; Rastas, 2017). Genotype data was converted to genotype likelihoods (posteriors) using the *linkage2post* script in Lep-Map2. Missing or erroneous parental genotypes were imputed using the *ParentCall2* module. SNP markers informative for both parents were assigned to linkage groups (LGs) using the *SeperateChromosomes2* algorithm in Lep-Map3 with lodLimit=11 and distortionLod=1. Unassigned SNPs were added to the preliminary map using the *JoinSingles2All* module with lodLimit=8, lodDifference=2, and distortionLod=1. The ordering of markers within LGs was conducted using the *OrderMarkers2* module after filtering markers based on segregation distortion (dataTolerance = 0.01). For each LG, the relative ordering of SNP markers was iterated ten times, and the configuration with the highest likelihood selected to represent a sex-averaged map for *O. edulis*. One large gap (>10cM) was identified and manually removed from the distal end of LG 10.

### Synteny and gene family expansion analyses

Gene level synteny was compared between *OE_Roslin_V1* and genome assemblies for a range of bivalve species using an orthogroup based approach. A list of putative one-to-one orthologues between *O. edulis* and assemblies for *C. gigas* (NCBI accession: GCF_902806645.1) (Peñaloza et al., 2021), *C. virginica* (GCF_002022765.2), and *P. maximus* (GCF_902652985.1) (Kenny et al., 2020) were generated using Orthofinder v.2.3.11 (Emms & Kelly, 2019). An independent *O. edulis* genome assembly generated by Boutet et al. (2022) (NCBI bioproject: PRJNA772088) was also included. The genomic coordinates of each gene in the one-to-one orthologue list for any two species under comparison was extracted and circos plots generated using the package Circlize 0.4.14 (Gu et al., 2014).

We inferred gene family expansions in *O. edulis* building on a published strategy (Regan et al., 2021). The start-point was all predicted proteins from the genome assemblies of 16 bivalve species, inclusive of *OE_Roslin_V1* (Supplementary Table 1). Longest isoforms for each protein were retained using AGAT v0.4.4 (Dainat DH, 2020). These sequences were used to generate orthogroups in Orthofinder v.2.3.11 (Emms & Kelly, 2019). FastTree (Price et al., 2010) was used to infer gene trees per orthogroup, which were compared against the rooted species tree by Orthofinder to infer duplications/losses using a duplication-loss-coalescent model (Emms & Kelly, 2019). Kinfin v1.0 (Laetsch & Blaxter, 2017a) was used to identify orthogroups that showed evidence for gene expansion in *O. edulis* compared to other bivalves (Regan et al,. 2021). Orthogroups showing evidence for gene expansions in *O. edulis* were first filtered for a fold change value >2.5 compared to the mean for all other bivalves. Fold-change is defined as the number of genes per orthogroup for *O. edulis* divided by the mean number of genes per orthogroup across all other bivalve species. Orthogroups meeting this filter, but with < 8/16 species (inclusive of *O. edulis*) represented in the tree, were further removed unless both *C. gigas* and *C. virginica* were present in the tree. Gene expansions in the remaining trees were classified as follows: i) orthogroups showing >3-fold mean expansion in gene copy number in all Ostreidae species (*O. edulis, C. gigas* and *C. virginica*) vs. other bivalves (i.e. potential ancestral Ostreidae expansion), plus a further >3-fold mean expansion in gene copy number comparing *O. edulis* to the mean for *C. gigas* and *C. virginica* (i.e. additional lineage-specific expansion in *Ostrea*), ii) orthogroups showing >3-fold mean expansion in gene copy number in all Ostreidae species, with no further expansion in gene copy number comparing *O. edulis* to the mean for *C. gigas* and *C. virginica* (i.e. inferred ancestral Ostreidae expansion only), iii) orthogroups showing >3-fold mean expansion in gene copy number in *O. edulis* vs. other bivalves, with no evidence for expansion in the Ostreidae ancestor (i.e. inferred lineage-specific expansion in *Ostrea* post-divergence from *Crassostrea*), iv) orthogroups showing >3-fold mean expansion in gene copy number in *O. edulis* compared to the mean for *C. gigas* and *C. virginica*, but lacking genes for other bivalve species (i.e. inferred Ostreidae specific genes showing lineage-specific expansion in *Ostrea* post-divergence from *Crassostrea*), v) orthogroups retaining genes for all three Ostreidae species, but lacking any genes for other bivalve species (i.e. inferred Ostreidae specific genes that have not shown further expansion) and vi) orthogroups showing >3-fold mean expansion in gene copy number in *O. edulis* compared to the mean for other non-Ostreidae bivalve species, absent in both *Crassostrea* species (inferred lineage-specific losses in *Crassostrea*, but lineage-specific expansion in *Ostrea*).

Functional annotation of each orthogroup was performed by searching each protein against the eukaryotic SignalP database (Petersen et al., 2011), Gene Ontology database (GO) (The Gene Ontology Consortium, 2019), and Pfam database (El-Gebali et al., 2019) using InterProScan v5.47-82.0 (Jones et al., 2014) (the top GO/Pfam/InterProScan annotation per orthogroup was recorded) and feeding the results into KinFin (Laetsch & Blaxter, 2017a). Functional annotations were summarised based on their counts across all the expanded orthogroups. Protein sequence alignments from selected orthogroups were retrieved and maximum-likelihood phylogenetic trees were generated using IQTREE v1.6.8 (Nguyen et al., 2015) using the best fitting substitution model (Kalyaanamoorthy et al., 2017) and running the ultrafast bootstrapping (Minh et al., 2013) for 1000 iterations to generate branch support value. The trees were then visualised using iTOL online server (Letunic & Bork, 2021).

## Results

### Contig assembly and quality evaluation

PromethION sequencing yielded 20,061,494 reads summing to 143.42 Gb of basecalled data with N50 length of 9,297 bp (Supplementary Figure 3) and mean length of 7,149 bp, which was used for contig assembly. Assuming a haploid genome size of 1.14 Gb following past flow cytometry work involving n=20 flat oysters sampled from Galicia in Spain (Rodríguez-Juíz et al., 1996), ∼120x long-read sequencing depth was achieved, including 26x with reads >15 Kb. Around 281 million Illumina short reads (∼72x sequencing depth) were used for genome polishing. Around 57.6 million Illumina reads were generated by sequencing the Omni-C™ library, which were used for genome scaffolding. RNA-Seq generated ∼50 million Illumina reads per tissue for genome annotation. K-mer based estimation predicted the *O. edulis* genome to be 881 Mb, with repeat content of 437 Mb (i.e. 49.8% of genome) and a heterozygosity rate of 1.02% (Supplementary Figure 4).

The Flye assemblies *OE_F1, OE_F2* and *OE_F3* were 976.2 Mb, 1,027.5 Mb and 964.2 Mb, respectively. Purging for haplotigs resulted in removal of 2-3% data across each assembly (Supplementary Table 2). The purged Flye assemblies had contig N50 values of 0.43, 0.39 and 0.34 Mb, respectively (Supplementary Table 2). Thus, OE_F1, which used a minimum overlap of 10,000 bp to generate a contig, had the highest contiguity. The wtdbg2 contig assembly *OE-RB1* was 829.1 Mb after purging and had an N50 value of 0.67 Mb (Supplementary Table 2). All four contig assemblies had a high BUSCO completeness score (∼90% complete) compared to the mollusca_odb10 database (Supplementary Table 2). The final merged and haplotig purged contig assembly *OE_contig_v1* was 934.9 Mb with a contig N50 of 2.38 Mb. Two rounds of genome polishing resulted in minor changes to contiguity, but increased BUSCO completeness from 89% to 95.2% (Supplementary Table 2), indicative of a strong positive effect on sequence accuracy.

### O. edulis chromosome level genome assembly

Scaffolding using HiRise and Juicer led to assemblies of 935.08 and 936.34 Mb with N50 values of 94.05 and 82.94 Mb, respectively (Supplementary Table 3). As the HiRise assembly was markedly more contiguous, it was taken forward as the basis for the final reference genome. Based on two lines of 3D contact evidence within the Omni-C data (see Methods), two large scaffolds in the HiRise assembly (scaffolds 11 and 12) were manually inserted into the super-scaffolds of the HiRise assembly. Specifically, scaffold 12 was inserted into super-scaffold 1 (at insertion point 65.4 Mb) and scaffold 11 was inserted at the start of super-scaffold 6 (Supplementary Figure 1). As noted in the methods, at this stage, super-scaffold 6 was renamed super-scaffold 2 as a product of it becoming the second largest scaffold in the HiRise assembly, maintaining the convention of naming scaffolds according to size (Supplementary Table 4).

The final assembly including the two manual corrections (*OE_Roslin_V1*) is 935.13 Mb with a scaffold-N50 of 95.56 Mb (Table 1), represented by 10 super-scaffolds comprising 93.65% (875.78 Mb) of the assembly, matching the haploid karyotype of *O. edulis* (i.e. 10 chromosomes) (Thiriot-Quiévreux, 1984; Leitao et al., 2002; Horváth et al., 2013). The remaining 59.3 Mb of *OE_Roslin_V1* comprises 1,353 unplaced scaffolds. The final assembly size matches closely to the k-mer based genome size estimate, and is slightly larger than other genome assemblies within Ostreidae, which could be due to lineage-specific repeat expansion (see later section).

**Table 1.** Genome statistics for *O. edulis* (*OE_Roslin_V1* assembly)

| **Metric** | **Value** |
| --- | --- |
| Assembly size (bp) | 935,138,052 |
| No. of contigs | 2,759 |
| Contig N50 (Mb) | 2.38 |
| Longest contig (Mb) | 16.06 |
| No of scaffolds | 1,363 |
| Length of top 10 scaffolds (bp) | 875,789,595 |
| Longest scaffold (bp) | 117,440,623 |
| Assembly N50 (bp) | 95,564,955 |
| Gaps (counts) | 1,534 |
| N's count | 153,250 |
| GC content (%) | 35.41 |
| Contigs > 500 bp | 1,363 |
| Contigs > 1000 bp | 1,294 |
| Contigs > 10,000 bp | 846 |
| Contigs > 100,000 bp | 103 |
| Contigs > 1Mb | 18 |

Detecting and correcting structural errors arising during genome assembly is critical in achieving a high-quality reference genome (Chen et al., 2021c). Evaluation of the assembly for structural errors identified 1,126 (663 expansions, 387 collapses, 76 inversions) putative structural errors when benchmarked against the raw nanopore reads, which were corrected. Assembly screening revealed little contamination from other taxa (Supplementary Figure 5). We observed a 97.09% mapping rate of nanopore reads back to the assembly, further demonstrating the accuracy and completeness of the reference genome. A K-mer copy number histogram revealed that haplotig purging was very efficient (Figure 1b). We identified 4,865 (91.9%) complete single copy BUSCO genes and 131 (2.5%) complete duplicated BUSCO genes in the final assembly (Figure 1c).

### Linkage map and assembly validation

The *de novo* variant calling pipeline called 24,522 SNPs across the ddRAD-Seq dataset. After stringent filtering (see Methods), the finished genetic map contained 4,016 SNPs anchored to the ten expected LGs (Supplementary Figure 6). We observed an overall high collinearity between these LGs and the *OE_Roslin_V1* genome assembly pseudo-chromosomes (Figure 1d, Supplementary Figure 7) confirming the accuracy of the scaffolding performed using the Omni-C data, including at the two manual joins we performed within the scaffold_1 and scaffold_2 of the *OE_Roslin_V1 assembly* (Figure 1d; Supplementary Figure 7). We observed a potential inversion between LG1 and super-scaffold 1, which was unrelated to the manually scaffolded region (Supplementary Figure 7). However, on closer inspection, the Hi-C data was ambiguous in this region (Figure 1a), with the opposite orientation of this region within the assembly being impossible to exclude, which would then match LG1.

### Genome annotation

57.3% (535.9 Mb) of the *OE_Roslin_V1* assembly was identified as repeats (Figure 2a), which falls in a similar range to recently published *C. gigas* genome assemblies (reported as 43% by Peñaloza et al. (2021) and 57.2% by Qi et al. (2021)). A large majority of repeats, comprising 37.65% of the genome, were annotated as unclassified (Figure 2a). A substantial proportion of the genome was annotated as LINE elements (5.98%), DNA transposons (4.37%) and rolling circles repeats (5.47%) (Figure 2a). The accompanying sister article to this study provides a more detailed curation of repeat landscape in an independently generated French *O. edulis* genome assembly (Boutet et al., 2022). Note, that this work identified a very similar proportion of repeats (55.1%) using the same bioinformatic pipeline, but not all could be confidently annotated.

**Figure 2.**
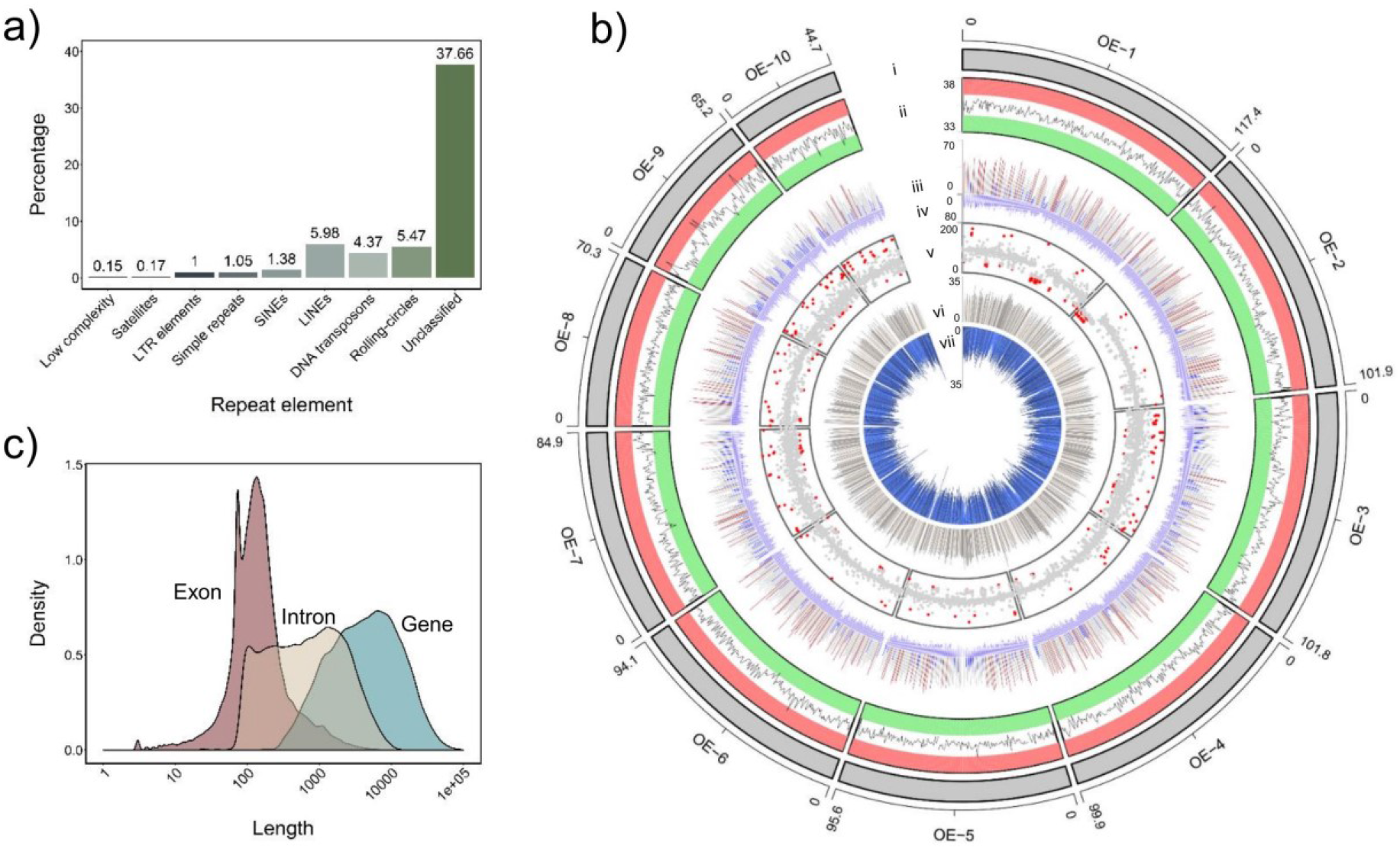
Annotation of the *O. edulis OE_Roslin_V1* assembly. a) Summary of genome repeat classes. b) Circos plot highlighting annotated features across the ten super-scaffolds (window size 0.5 Mb except track-v, which is 0.1Mb). Tracks as follows: i: 10 super-scaffolds OE-1 to OE-10, ii: GC percentage (33-38%), with red and green bars indicating GC >36.5% and < 34.5%, respectively, iii: Genic content (sum of annotated gene models) expressed as percentage of total window size, regions with <20% genic content are coloured blue, while 20 to 40% are coloured grey and >40% are coloured red, iv: Gene density (0-80). v: mean Illumina sequencing depth, with values < 45 and > 150 shown as red points, vi: classified repeats expressed as percentage of total window size (0 to 35%), vii: Novel unclassified repeat elements expressed as percentage of total window size (0 to 35%), c) Density plot showing gene, exon and intron lengths.

Gene model prediction identified 35,699 coding genes in the masked genome (Table 2). Genic regions comprised 261.83 Mb (28.42%) of the genome size, with an average gene length of 7,411 bp (Figure 2c) and an average coding sequence length of 1,224 bp. Functional annotation of the predicted proteins resulted in annotation of 23,109 gene models with EggNOG hits and provided 17,504 gene models with a GO annotation (Table 2). A range of annotate features are plotted along the genome in Figure 2b.

**Table 2.** Genome annotation statistics for *O. edulis* (*OE_Roslin_V1*)

| **Metric** | **Value** |
| --- | --- |
| Protein coding genes | 35,699 |
| Average gene length (bp) | 7,411 |
| Average exon length (bp) | 241 |
| Single exon transcripts | 1,631 |
| Multiple exon transcripts | 34,068 |
| Total gene length (bp) | 265,862,173 |
| **Functional annotation (No of proteins)** |  |
| GO annotation | 17,504 |
| Interproscan hits | 19,613 |
| Eggnog hits | 23,109 |
| Pfam hits | 16,966 |
| Cazyme hits | 537 |
| Merops hits | 921 |

### Additional validation of manually incorporated scaffolds

To confirm the validity of the manually scaffolded regions in super-scaffolds 1 and 2, we sought to concretely demonstrate that they belonged to the flat oyster genome. We firstly performed BLASTn (Altschul et al., 1997) searches for all coding genes predicted in these regions against *C. gigas* (Peñaloza et al. 2021) and an independent *O. edulis* assembly (Boutet et al. 2022), and compared the results to the remaining regions of super-scaffolds 1 and 2 (summarized in Supplementary Table 5; raw data in Supplementary Table 6). The proportion and percentage identity of BLAST hits to both oyster genomes was highly comparable for both regions along super-scaffolds 1 and 2. Secondly, RNA-Seq reads (pooled from heart, striated muscle and gonad) mapped with variable depth to approximately 40% of the predicted genes within the manually incorporated regions of super-scaffold 1 and 2 (Supplementary Figure 8). The RNA-Seq mapping rate and depth was lower in the manually incorporated regions than the remaining parts of super-scaffolds 1 and 2 (Supplementary Figure 8).

### Synteny analysis with other bivalve genomes

Synteny plots of 1-to-1 orthologue gene locations revealed conserved chromosomal-level synteny between *OE_Roslin_V1* and three independently assembled bivalve genomes: *C. gigas* (Figure 3a), *C. virginica* (Figure 3b) and *P. maximus* (Figure 3c). We observed little evidence for major chromosomal rearrangements (i.e. involving megabases of a chromosome undergoing inversion or translocations) between the 10 chromosomes of *O. edulis* and *C. gigas* (Figure 3a), indicating that the ancestral ostreid karyotype has been maintained in both species. Comparison of *OE_Roslin_V1* with *C. virginica* (Figure 3b) provides evidence for possible chromosomal rearrangements in *C. virginica* after its split with *C. gigas*, assuming the chromosome-level synteny between *O. edulis* and *C. gigas* reflects the ancestral state. For instance, super-scaffold 8 in *OE_Roslin_V1*, which shares synteny across the length of *C. gigas* chromosome 4, shares synteny with two major blocks on *C. virginica* chromosomes 5 and 6 (Figure 3b). The synteny relationship between *OE_Roslin_V1* and the extensively rearranged *P. maximus* genome was consistent with that reported between *C. gigas* and *P. maximus* (Yang et al., 2021). We observed genome-wide synteny between *OE_Roslin_V1* and an independently generated assembly for *O. edulis* (Boutet et al. 2022), although there were a small number of chromosomal regions where synteny was broken (Figure 3d).

**Figure 3.**
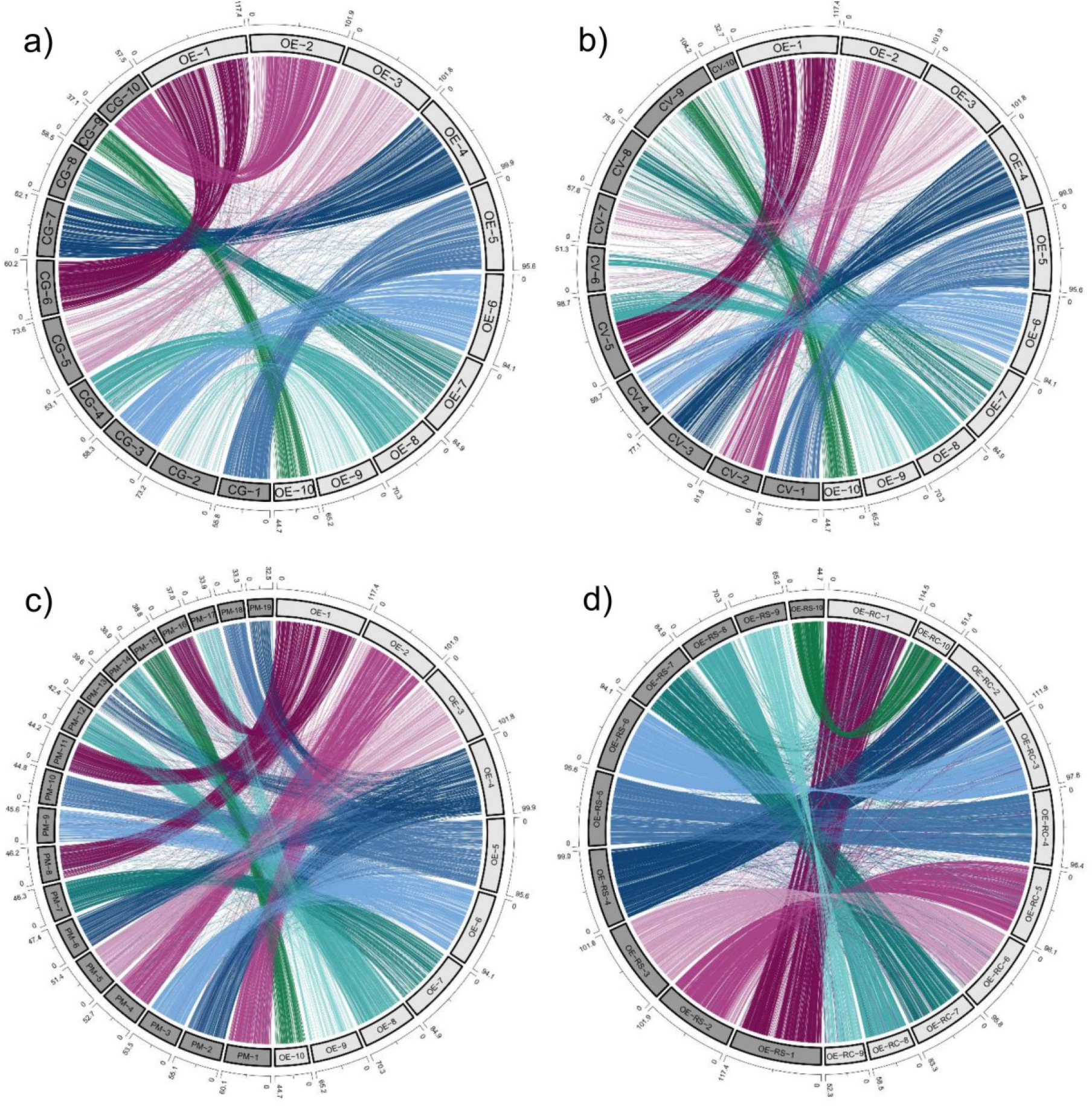
Chromosome level synteny between the *OE_Roslin_V1 O. edulis* assembly and three independent bivalve assemblies. Circos plots are shown comparing the ten super-scaffolds (OE1-OE10) with putative chromosomes of a) *C. gigas*, b) *C. virginica*, c) *P. maximus* chromosomes, and d) an independent *O. edulis* assembly reported in Boutet et al. (2022) (‘RC’ denotes super-scaffolds from Boutet et al. (2022); ‘RS’ denotes super-scaffolds from *OE_Roslin_V1*).

### Gene families expanded during Ostrea evolution

Gene duplication is associated with adaptation during evolution (Ohno, 1970), including in bivalves (Phuangphong et al., 2021; Regan et al., 2021). To gain insights into how gene duplication influenced *Ostrea* evolution, we identified gene family expansions in *OE_Roslin_V1* by comparison to 15 additional bivalve genomes. 712 gene families showed evidence of expansion (Supplementary Table 7; see Methods), categorized into six groups in a phylogenetic framework (Figure 4a). The most common class of putative gene family expansion involved genes distributed among different bivalve families that underwent expansion in Ostreidae (Figure 4b), with a subset showing evidence of further expansion in *O. edulis* compared with the two *Crassostrea* species (Figure 4c). Similarly, we observed many gene families distributed among several bivalve families, where expansion was specific to *Ostrea* (Figure 4d). We also identified gene families specific to all three Ostreidae members (i.e. absent in other bivalves), among which a large proportion did not show further expansion in *O. edulis* compared to *Crassostrea* (Figure 4e), with a smaller group expanded in *O. edulis* specifically (Figure 4f). Finally, we found a small number of gene families represented by different bivalve families that showed expansion in *O. edulis*, but absence in *Crassostrea* species (Figure 4g).

**Figure 4.**
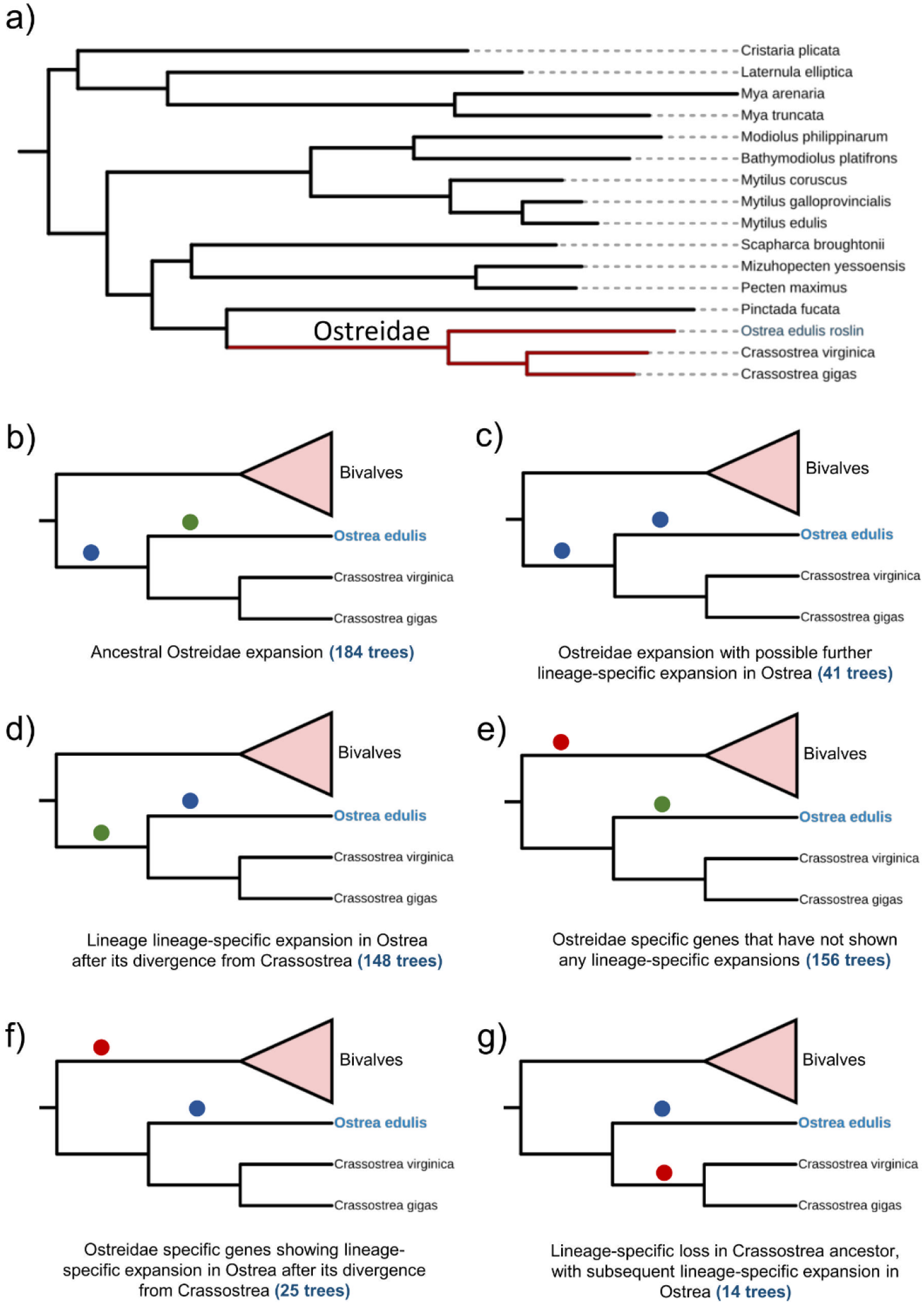
Classification of gene family expansion during *O. edulis* evolution. a) Species tree of bivalve genomes used in the analysis, b-f) different categories of gene family expansion (classified as described in Methods). Branch annotations: Blue circles indicate putative expansion; Green circles indicates no expansion; Red circle indicates an absence of species along that branch for the affected orthogroups. Full data is provided in Supplementary Table 7.

Annotation of protein domains in the expanded gene families may offer clues into biological functions targeted during *Ostrea* evolution (Supplementary Table 7; summarized in Figure 5a). Among 701 expanded gene families annotated with conserved domains by Interproscan (Jones et al., 2014), 229 were unique to 1 gene family, with the remaining domains present in 2 to 31 gene families. Thus, many domains were overrepresented among the expanded gene families (Figure 5a), including G protein-coupled receptor, rhodopsin-like (IPR000276; 31 gene families) and secretin-like (IPR000832; 9 gene families). Several domains associated with innate immune function were overrepresented, including C-type lectin (IPR001304; 20 gene families), complement C1q (IPR001073; 15 gene families), and Sushi/SCR/CCP (i.e. complement control protein domain) (IPR000436; 9 gene families). There were many overrepresented domains containing zinc finger motifs (including IPR000315; 18 gene families, IPR013087; 9 gene families; and IPR001878; 5 gene families). The highly conserved homeobox domain was annotated in 6 gene families expanded in *O. edulis*. We provide two examples of expanded gene families in Figure 5b and c, both OGs taken from gene families showing lineage-specific expansion in *Ostrea* after its divergence from Crassostrea.

**Figure 5.**
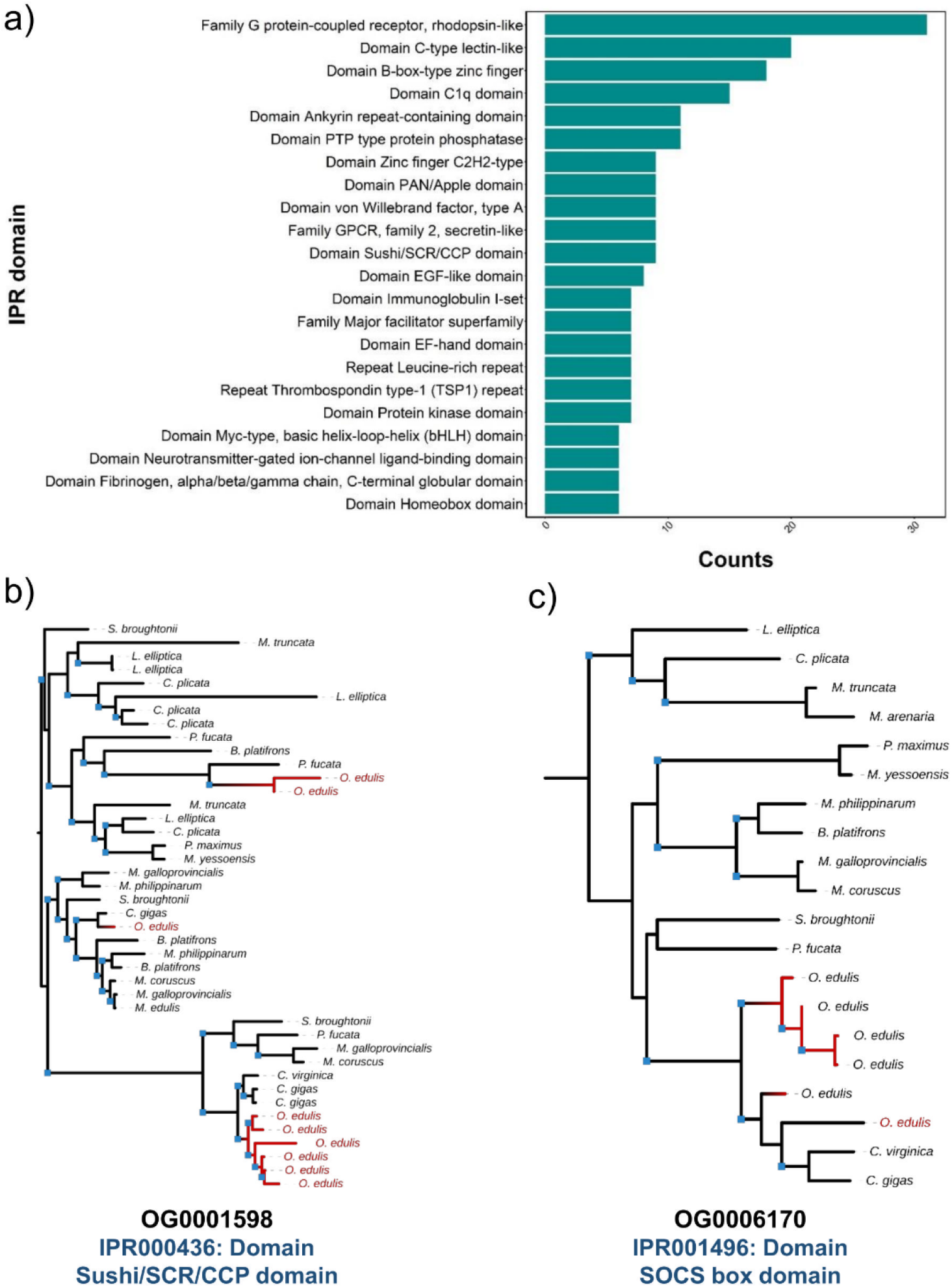
Most represented protein domains in expanded *O. edulis* gene families. a) Top 20 represented IPR domains. b) & c) Example maximum likelihood phylogenetic trees highlighting gene family expansions in *O. edulis*. Blue squares at nodes indicate bootstrap support value >50%.

We further used this dataset to identify extremely expanded gene families in the *O. edulis* genome. For instance, we observed two orthogroups showing massive tandem expansion of genes encoding proteins with the uncharacterized EB domain (IPR006149). In both cases, these gene families were specific to Ostreidae and present as either 1 or 2 copies in *Crassostrea* species, but 31 (orthogroup OG0002210) and 11 copies (orthogroup OG0013280) in *O. edulis* (Supplementary Table 7). There were many other gene families specifically highly expanded in *O. edulis* (Supplementary Table 7), including an Ostreidae specific family (orthogroup OG0001484) encoding proteins containing a SAP domain (41 genes in *O. edulis*, vs. 2 genes each in both *Crassostrea* species), which has been proposed to be involved in chromosomal organization (Aravind & Koonin, 2000).

## Discussion

The high-quality, publicly available genome assembly we have generated and annotated for *O. edulis* serves as a novel reference for genetics investigations of wild and farmed European flat oyster, in addition to comparative genomic investigations of molluscan taxa. Additional resources of value to the research community have been produced and made publicly available, including multi-organ RNA-Seq data, which we used to support gene model prediction and confirm genome assembly quality, but in the future can be used to explore patterns of tissue gene expression. In terms of assembly quality, the contig N50 we achieved is among the highest of all bivalve assemblies publicly available. This demonstrates the utility of our choice to merge different contig assemblies using Quickmerge (Chakraborty et al., 2016), which has been shown elsewhere to be effective for generating high-quality assemblies in molluscs (Sun et al., 2021), and other taxa (e.g. Chen et al., 2021a; Li et al., 2021; Mathers et al., 2021). Genome-wide sequence accuracy was further evidenced by the high mapping rate of nanopore reads back to the assembly, and the limited number of structural errors in the genome, which was lower than reported for the recent *C. gigas* reference genome (Peñaloza et al. 2021). BUSCO scores for our final *O. edulis* assembly are in the range of high-quality molluscan genome assemblies published to date (e.g. Sun et al, 2021), indicating an excellent level of gene representation.

Interestingly, our k-mer based genome size estimate (881 Mb), which matched closely with our final assembly length (876 Mb), was only ∼ 77% of the 1.14 Gb genome size previously estimated by flow cytometry in a population of Spanish flat oysters (Rodríguez-Juíz et al., 1996). Similar observations have been made for other bivalve genomes, including *C. gigas* (e.g. Peñaloza et al., 2021). The discrepancy between this past flow cytometry assessment and our own sequencing-based estimates could be partly explained by population differences in genome size, considering the plasticity of genome content within bivalve species (Gerdol et al., 2020). However, this discrepancy cannot be easily explained by an under-representation of repeats in our assembly, considering that >97% of the raw nanopore reads mapped back to the final assembly. Underestimation of genome size can also arise due to high heterozygosity (Liu et al., 2020). Our heterozygosity rate estimate of 1.02% for *O. edulis* was within the range reported for other bivalves, including 1.3% in *C. gigas* (Zhang et al., 2012) and 1.04% in scallop (*Patinopecten yessoensis*) (Wang et al., 2017). This is interesting, as these previous estimates were made using individuals selected for reduced heterozygosity via inbreeding (Zhang et al., 2012) or by using a selfing family (Wang et al., 2017), implying a possible loss of genetic diversity in the *O. edulis* population we used for sequencing (e.g. a historic bottleneck). In contrast, an outbred *C. gigas* individual recently sequenced showed a much higher heterozygosity rate estimate of 3.2% (Peñaloza et al., 2021).

With regards to genome annotation, the average gene length we obtained (7,411 bp; Figure 2c) is lower than high-quality annotations for oyster genome assemblies, for example the *C. gigas* reference genome annotated by NCBI RefSeq (PRJNA629593) has almost twice the average gene length (10,990 bp). Considering the high accuracy, completeness and contiguity of our assembly, the result cannot be explained by differences in assembly quality. Instead, it is likely that our annotation strategy was inefficient in predicting gene models compared to NCBI RefSeq, leading to more fragmented or partially predicted gene models, explaining the reduced length statistics. However, our annotation still has global utility, considering that we observe extensive 1-to-1 orthologue mapping compared to other genome assemblies (Figure 3), and were able to perform valid comparative genomic analyses both here (i.e. Figure 4, 5) and in studies that have used our annotation to date (see later paragraph). The reader should also be aware that our assembly will undergo NCBI RefSeq annotation in the near future, which will improve the quality of gene prediction, in turn enhancing future genetics and comparative genomic investigations exploiting the genome as a reference. In the longer-term, we anticipate that bivalve genomes will benefit from greatly improved functional annotations that extend far beyond gene model prediction, incorporating functional assays defined by the FAANG initiative to identify chromatin state modifications, regulatory elements, non-coding RNAs and isoform diversity (Clark et al., 2020).

Our cross-species synteny analysis revealed few major chromosomal reorganisations in the flat oyster genome, consistent with previous reports describing the near conserved karyotype across all oysters (Guo et al., 2018). Furthermore, conserved synteny and chromosomal architecture against an independently assembled flat oyster genome assembly (Boutet et al., 2022), coupled with the general high congruency of the assembled super-scaffolds with linkage groups, further confirmed the global quality of our assembly. Expansions to gene families involved in stress responses during bivalve evolution may reflect adaptation to a filter-feeding sessile lifestyle in a hostile environment (Guo et al., 2018; Regan et al., 2021; Hu et al., 2022). Past work has revealed expansions in gene families encoding heat shock proteins, as well as families involved in apoptosis inhibition and innate immunity, including C-type lectins and C1q complement domain containing proteins. The gene family expansions reported here mirror these adaptation strategies, with enrichment in functional annotations for pathogen recognition and inflammatory response, e.g. C type lectins, complement and immunoglobulin domains. The comparative genomic resources provided here can support future evolutionary analyses of gene families, and should prove useful when interpreting the fine mapping of genetic variation around flat oyster genes, for instance those identified in QTL regions.

Future applications of the *O. edulis* reference genome reported here, and for an independent genome assembly described for a French *O. edulis* individual in an accompanying article (Boutet et al., 2022) will address challenges relating to flat oyster conservation and sustainable aquaculture production. These genomes provide researchers with new tools that empower genetic approaches addressing the ubiquitous threat posed by *Bonamia* via a range of technologies (Houston et al., 2020; Potts et al., 2021). In this regard, the genome reported here is proving useful already, with a recent study revealing that SNP markers previously associated with *Bonamia* resistance (Vera et al., 2019) are located in high linkage-disequilibrium across a large region of super-scaffold 8, which contains many candidate immune genes (Martinez et al. 2022). Another recent study from has mapped variants genotyped with an existing medium density SNP array (Gutierrez et al., 2017) against our new *O. edulis* genome, identifying QTLs underpinning variation in growth traits on super-scaffold 4 (Peñaloza et al., 2022). Via its public release with all accompanying raw data, we anticipate rapid uptake of our genome by the research community, and envisage the next steps for the field to include broader surveys of genome-wide diversity covering a global representation of populations. This new phase of genome enabled biology is like to uncover many secrets on the genetic and functional basis for adaptation and disease resilience in this iconic oyster species.

## Acknowledgements

This study was funded by the Biotechnology and Biological Sciences Research Council (BBSRC) under the AquaLeap consortium (grant code: BB/S004181/1) and received additional support from BBSRC Institute Strategic Programme grant BBS/E/D/10002070. We thank Edinburgh Genomics, especially Marian Thomson, for performing the PromethION sequencing and providing associated advice leading up to the work.

## Author contributions

MKG, RDH, TPB and DJM conceptualized the study. MKG sampled the sequenced oyster, extracted DNA and RNA used for sequencing, and led the genome assembly and annotation. IB and AT performed lab work and generated the ddRAD-Seq data for linkage map construction. CP led the linkage map construction. TR and MKG performed the gene-family expansion analysis. MKG and DJM co-wrote the manuscript with inputs from all authors leading to the submitted manuscript.

