## Supplementary data for "Chromosome level reference genome for European flat oyster (*Ostrea edulis* L.)"

**Supplementary Table 1**. Details of species used in the gene family expansion analysis

| **Species** | **Predicted proteins** | **Proteins assigned to orthogroups** | **Data source** | **Data URL** |
| --- | --- | --- | --- | --- |
| *Bathymodiolus platifrons* | 33,581 | 32,344 | MDB | <https://download.molluscdb.org/v2/> |
| *Crassostrea gigas* | 31,371 | 30,767 | NCBI | <https://ftp.ncbi.nlm.nih.gov/genomes/all/GCF/902/806/645/GCF_902806645.1_cgigas_uk_roslin_v1/GCF_902806645.1_cgigas_uk_roslin_v1_protein.faa.gz> |
| *Crassostrea virginica* | 34,587 | 33,824 | NCBI | <https://ftp.ncbi.nlm.nih.gov/genomes/all/GCF/002/022/765/GCF_002022765.2_C_virginica-3.0/GCF_002022765.2_C_virginica-3.0_protein.faa.gz> |
| *Cristaria plicata* | 30,047 | 25,022 | MDB | <https://download.molluscdb.org/v2/> |
| *Laternula elliptica* | 36,686 | 29,048 | MDB | <https://download.molluscdb.org/v2/> |
| *Mizuhopecten yessoensis* | 24,515 | 23,040 | NCBI | <https://ftp.ncbi.nlm.nih.gov/genomes/all/GCF/002/113/885/GCF_002113885.1_ASM211388v2/GCF_002113885.1_ASM211388v2_protein.faa.gz> |
| *Modiolus philippinarum* | 36,549 | 35,525 | MDB | <https://download.molluscdb.org/v2/> |
| *Mya arenaria* | 24,368 | 21,767 | MDB | <https://download.molluscdb.org/v2/> |
| *Mya truncata* | 48,080 | 42,443 | MDB | <https://download.molluscdb.org/v2/> |
| *Mytilus coruscus* | 58,249 | 55,155 | NCBI | <https://ftp.ncbi.nlm.nih.gov/genomes/all/GCA/011/752/425/GCA_011752425.2_MCOR1.1/GCA_011752425.2_MCOR1.1_protein.faa.gz> |
| *Mytilus edulis* | 60,330 | 56,691 | MDB | <https://download.molluscdb.org/v2/> |
| *Mytilus galloprovincialis* | 38,439 | 36,589 | MDB | <https://download.molluscdb.org/v2/> |
| *Ostrea edulis* | 35,699 | 31,617 | Current study | ---- |
| *Pecten maximus* | 26,122 | 24,191 | NCBI | <https://ftp.ncbi.nlm.nih.gov/genomes/all/GCF/902/652/985/GCF_902652985.1_xPecMax1.1/GCF_902652985.1_xPecMax1.1_protein.faa.gz> |
| *Pinctada fucata* | 31,477 | 29,231 | OIST Marine Genomics Unit | <https://marinegenomics.oist.jp/pearl/viewer?project_id=36> |
| *Scapharca broughtonii* | 24,045 | 21,906 | GigaDB | <ftp://parrot.genomics.cn/gigadb/pub/10.5524/100001_101000/100607/EVM.final.gene.gff3.pep> |

**Supplementary Table 2**. Statistics for various contig assemblies generated in the study

| Assembly ID | Assembly length pre-purging (Mb) | Assembly length (Mb) | No of sequences | N50 (Mb) | Purged content (% genome) | BUSCO complete  [C (S,D);F;M]% |
| --- | --- | --- | --- | --- | --- | --- |
| OE_F1_purged | 976.27 | 954.62 | 10,384 | 0.430 | 21.65 | [91.8(90.5:1.3);1.6;6.6] |
| OE_F2_purged | 1027.57 | 1001.95 | 9,961 | 0.398 | 25.62 | [89.4.1(86:3.4);1.4;9.2] |
| OE_F3_purged | 964.21 | 944.23 | 11,336 | 0.348 | 19.98 | [91.4(90:1.4);1.7;6.9] |
| OE_RB1_purged | 858.75 | 829.13 | 8,818 | 0.671 | 29.62 | [88.6(87.8:0.8);1.4;10] |
| OE_contig_v1 | -- | 933.6 | 2,759 | 2.38 | -- | [89(85.7:3.3);1.5;9.2] |
| OE_contig_pilon_v1 | -- | 934.9 | 2,759 | 2.38 | -- | [95.2(91.2:4);0.5;4.3] |

**Supplementary Table 3**. Assembly statistics for the Dovetail HiRise and Juicer assembly

| **Statistic** | **HiRise** | **Juicer** |
| --- | --- | --- |
| Assembly size (Mb) | 935.13 | 936.34 |
| No of scaffolds | 1,365 | 3,148 |
| N50 (Mb) | 94.05 | 82.94 |
| Largest scaffold (Mb) | 108.01 | 135.03 |
| Gaps | 1,531 | 2,809 |
| N’s counts (bp) | 153,100 | 1,364,100 |
| Length of top 10 scaffolds (Mb) | 852.78 | 813.53 |
| Length of top 20 scaffolds (Mb) | 889.34 | 845.80 |
| Length of scaffolds >10 Mb | 866.25 | 825.17 |
| No of scaffolds >5Mb | 12 | 11 |

**Supplementary Table 4.** Assembly and annotation statistics for the 10 Linkage groups

| Linkage group | Scaffold ID | Length (bp) | Protein coding genes | Percentage repeats |
| --- | --- | --- | --- | --- |
| LG01 | Scaffold_1 | 117,440,623 | 4,750 | 55.73 |
| LG03 | Scaffold_2 | 101,867,661 | 3,960 | 53.57 |
| LG08 | Scaffold_3 | 101,833,125 | 3,720 | 58.85 |
| LG02 | Scaffold_4 | 99,930,069 | 4,022 | 54.98 |
| LG05 | Scaffold_5 | 95,564,955 | 3,720 | 55.40 |
| LG06 | Scaffold_6 | 94,056,450 | 3,743 | 54.27 |
| LG04 | Scaffold_7 | 84,932,467 | 3,167 | 58.83 |
| LG07 | Scaffold_8 | 70,328,625 | 2,714 | 54.25 |
| LG09 | Scaffold_9 | 65,180,066 | 2,512 | 59.74 |
| LG10 | Scaffold_10 | 44,655,554 | 1,520 | 56.84 |

**Supplementary Table 5**. Summary of BLASTn outputs (<1e-20) comparing manually integrated and remaining regions of super-scaffolds 1 and 2.

| **BLAST database** | **Super-scaffold (region)** | **Genes annotated** | **Number hits vs. database** | **% genes with hit** | **Mean / SD % nucleotide identity vs. top hit** |
| --- | --- | --- | --- | --- | --- |
| Pacific oyster |  |  |  |  |  |
|  | 1 (manually integrated) | 466 | 57 | 12.23 | 85.92 / 4.31 |
|  | 1 (remaining region) | 5,604 | 686 | 12.24 | 84.81 / 4.64 |
|  | 2 (manually integrated) | 689 | 93 | 13.5 | 84.50 / 5.15 |
|  | 2 (remaining region) | 4,361 | 759 | 17.4 | 85.35 / 4.84 |
| Flat oyster |  |  |  |  |  |
|  | 1 (manually integrated) | 466 | 342 | 73.39 | 99.33 / 1.33 |
|  | 1 (remaining region) | 5,604 | 2585 | 46.13 | 99.08 / 2.12 |
|  | 2 (manually integrated) | 689 | 500 | 72.57 | 99.40 / 1.24 |
|  | 2 (remaining region) | 4,361 | 3,392 | 77.78 | 99.05 / 2.12 |

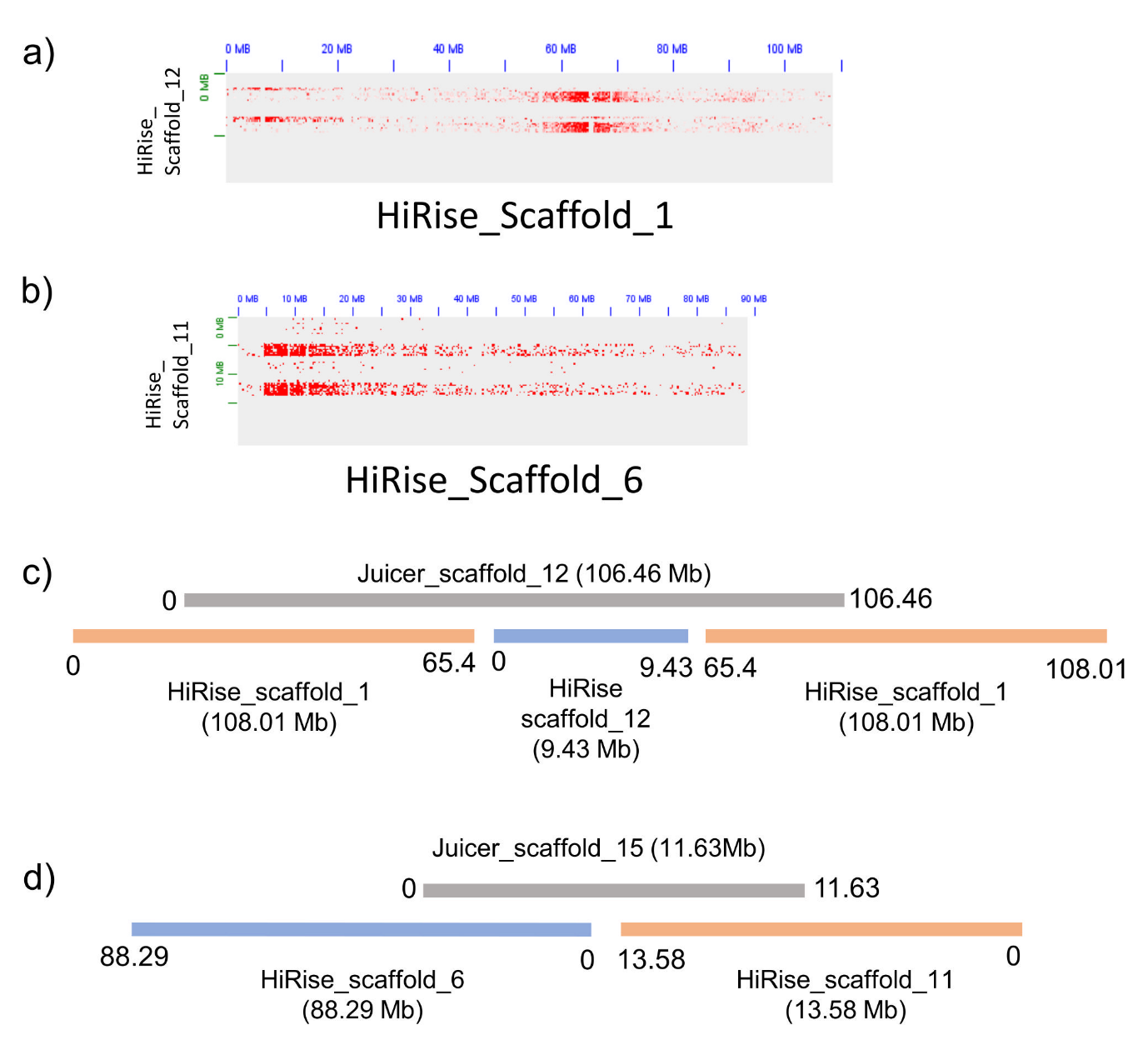

**Supplementary Figure 1**. a) and b) 3D interaction signals confirming interactions between Scaffolds 1 and 6 of the HiRise assembly with Scaffolds 12 and 11, respectively. c) and d) Representative visualisation of the alignments between the HiRise and Juicer assembly scaffolds which were used to confirm and manually join the scaffolds as observed in a) and b).

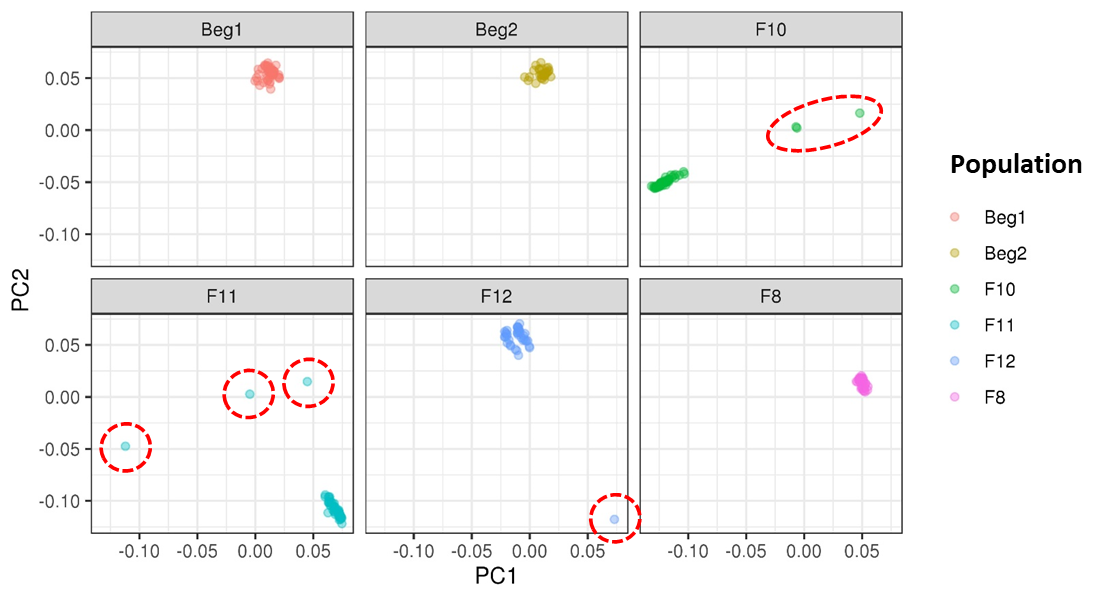

**Supplementary Figure 2**. PCA plot highlighting outliers detected across different oyster families used to construct the linkage map. Only families F10, F11, F12 and F8 were used for the construction of the linkage map.

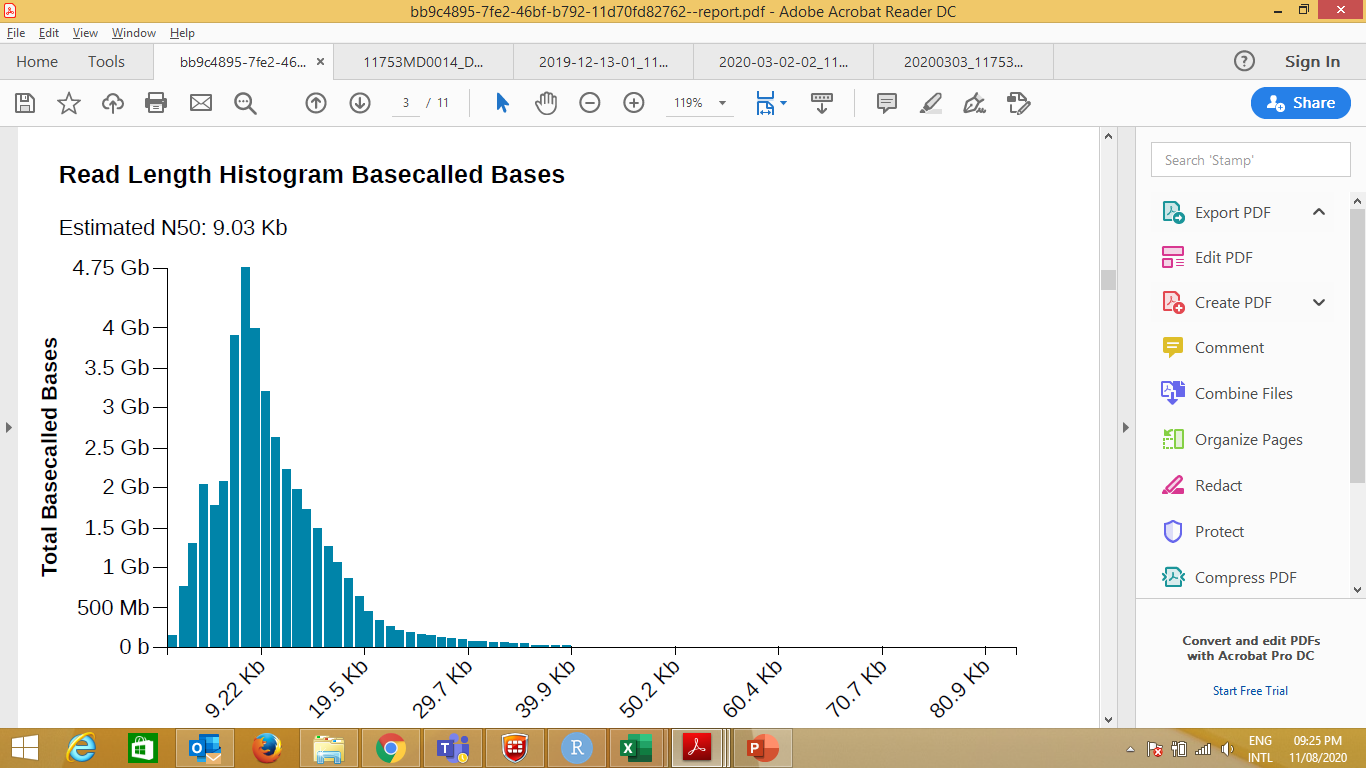

**Supplementary Figure 3.** Read length histogram of the raw Nanopore data

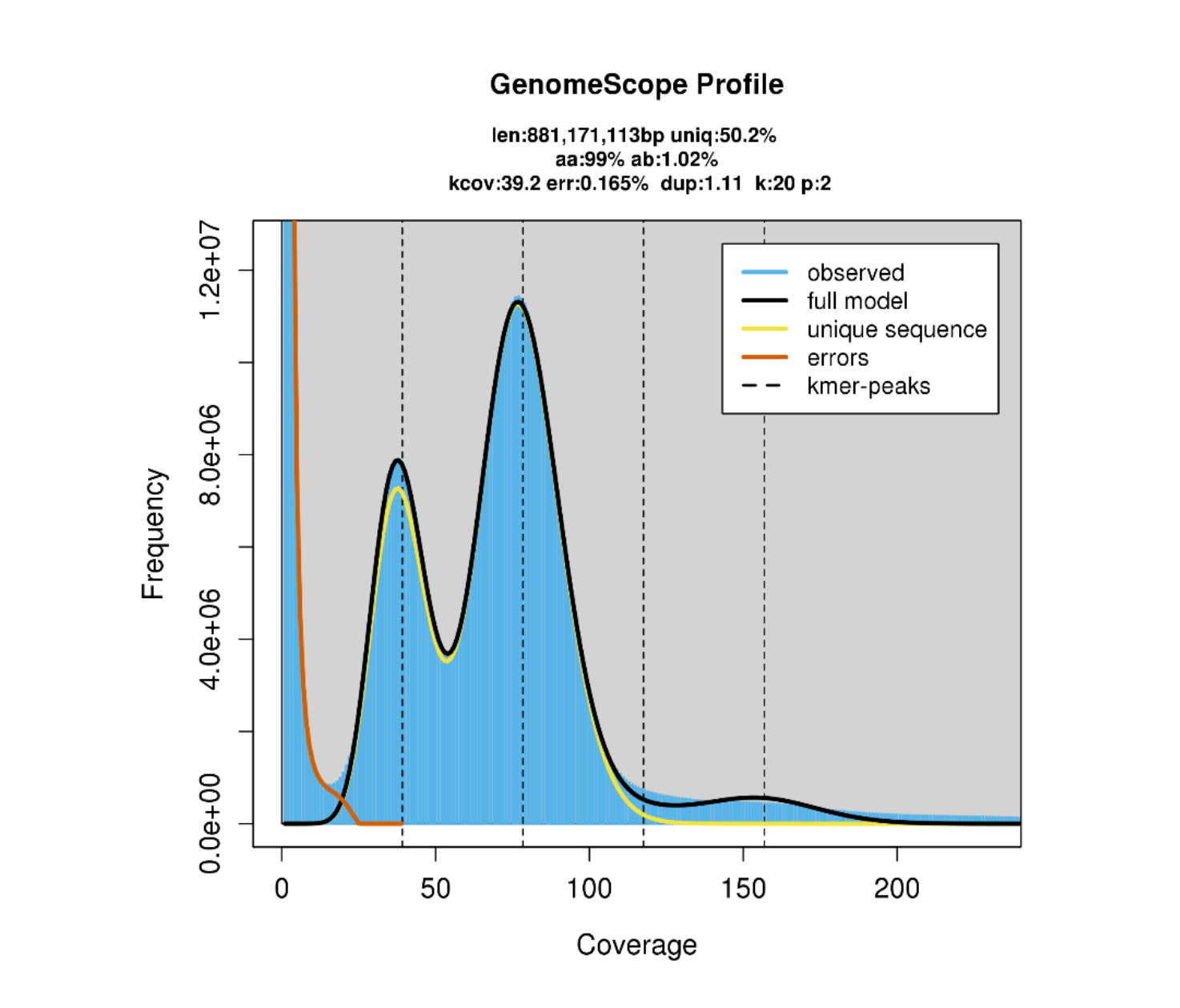

**Supplementary Figure 4**. Genomescope profile generated using a k-mer value of 20. Histogram highlights the k-mer profile of the *O*. *edulis* short read dataset. Blue curve indicates the observed k-mer frequencies and black line highlights the Genomescope model. We observe the error k-mers at low frequencies. The plot also displays the predicted genome size (“len”), heterozygosity (“ab”) and non-repeat percentage (“uniq”).

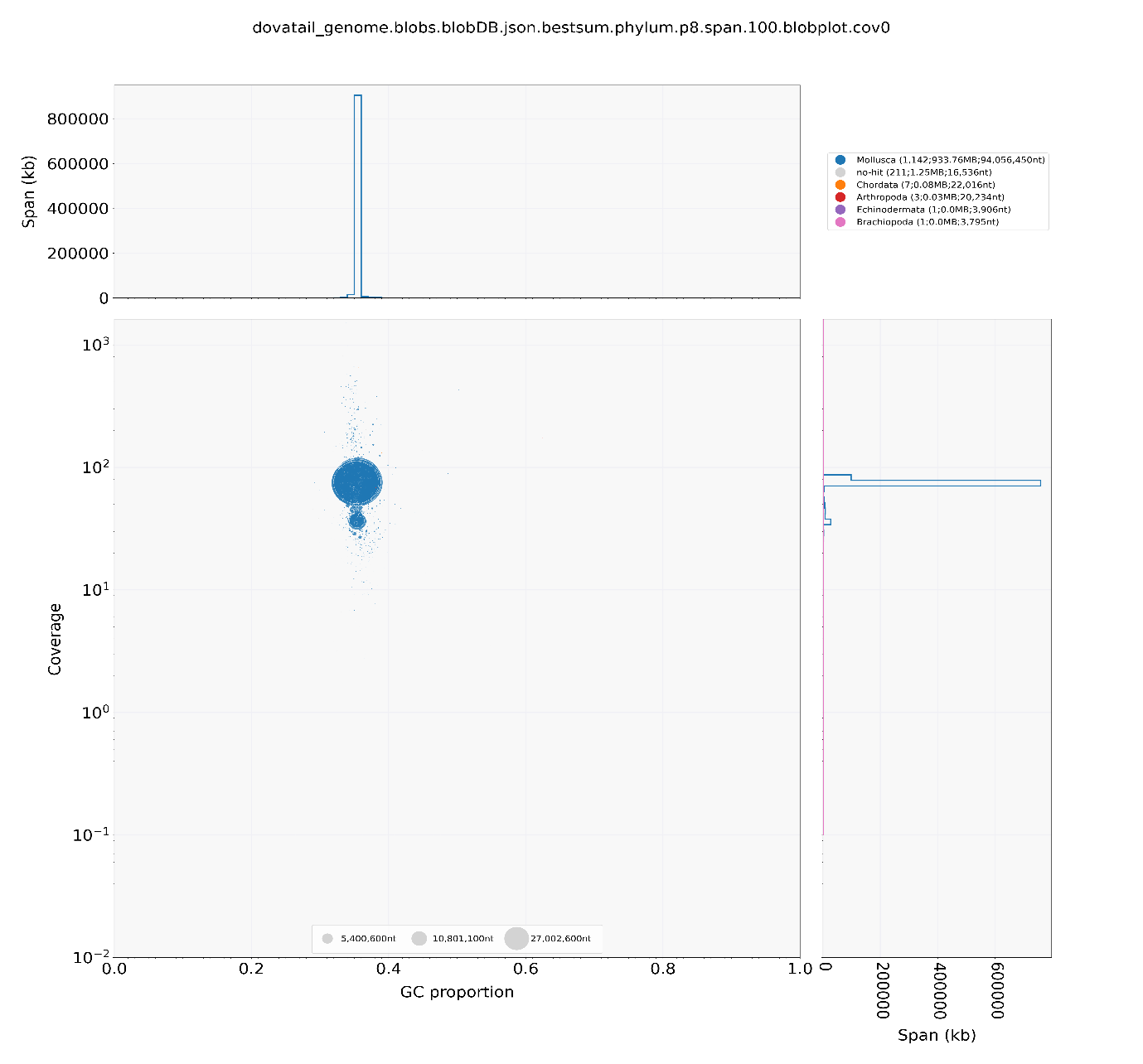

**Supplementary Figure 5**. Blobplot highlighting limited contamination within the *O. edulis* *OE_Roslin_V1* assembly with <0.01 Mb of all reads matching to phylum Chordata, Arthropoda, Echinodermata and Brachiopoda and no prokaryotic contamination. The coverage and GC content of these sequences match well with the coverage of the *O. edulis* sequence, it is therefore likely to be *O. edulis* specific sequences that are not in the NCBI database, but have a best hit against other Eukaryotic sequences.

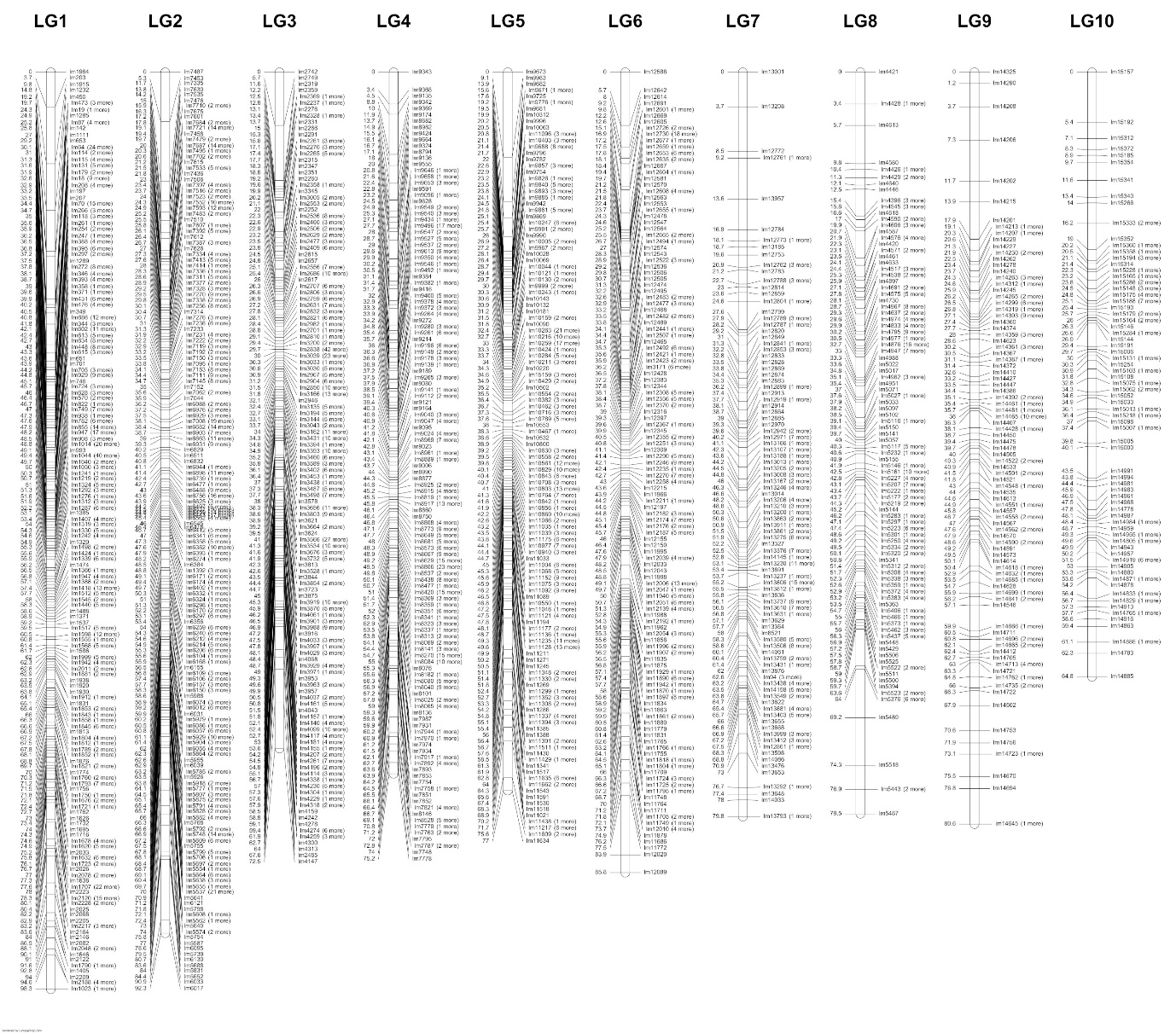

**Supplementary Figure 6**. Linkage map containing SNP markers aligning to the *O. edulis OE_Roslin_V1* assembly, visualized with LinkageMapView (Ouellette et al. 2017).

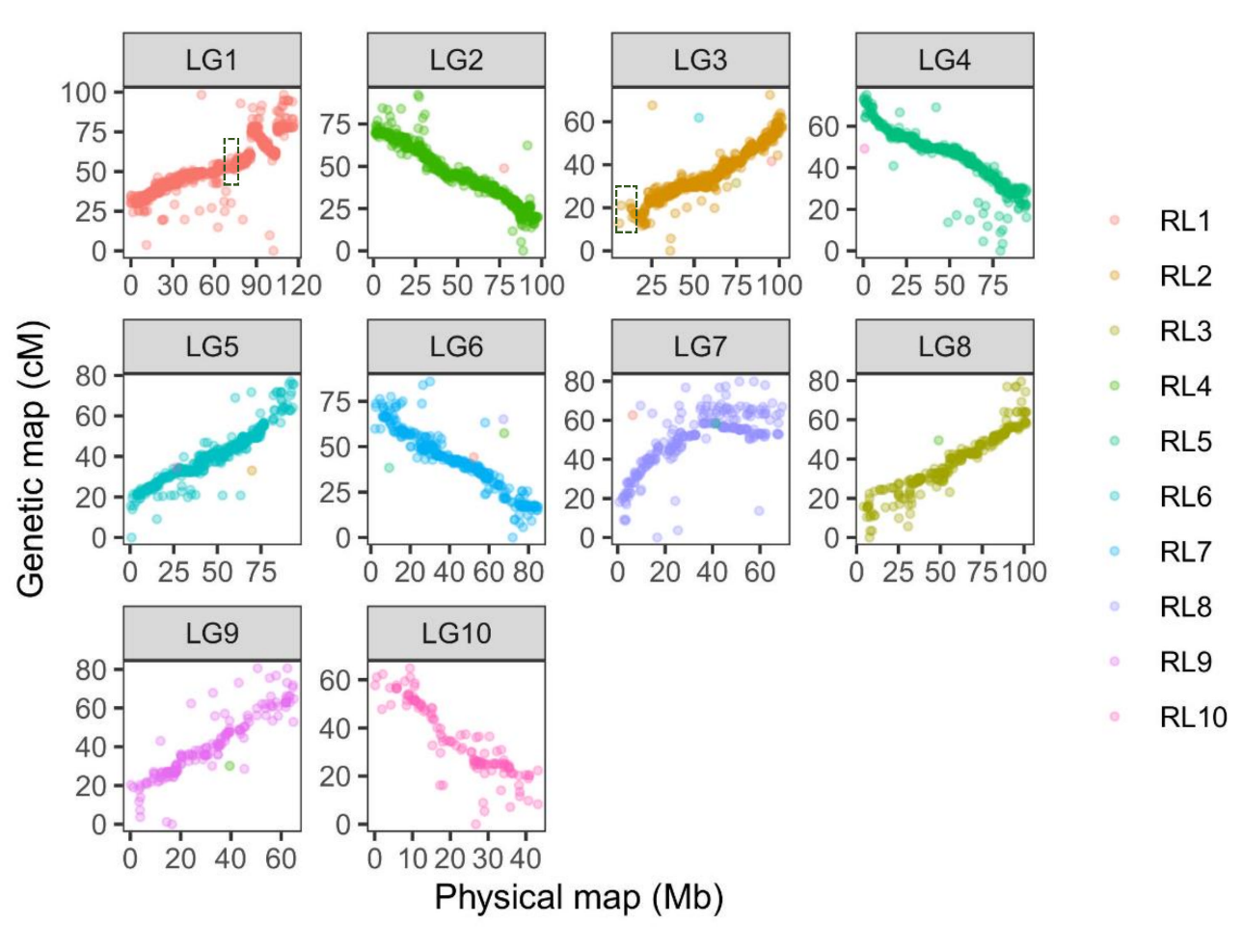

**Supplementary Figure 7**. A novel linkage map for *O. edulis* shows a high level of collinearity with the *OE_Roslin_V1* assembly. Each point represents a SNP in the linkage map that is coloured according to the super-scaffold to which it maps. The black squares within the linkage groups LG1 (super-scaffold 1) and LG3 (super-scaffold 2) indicate the regions where manual scaffolding was performed.

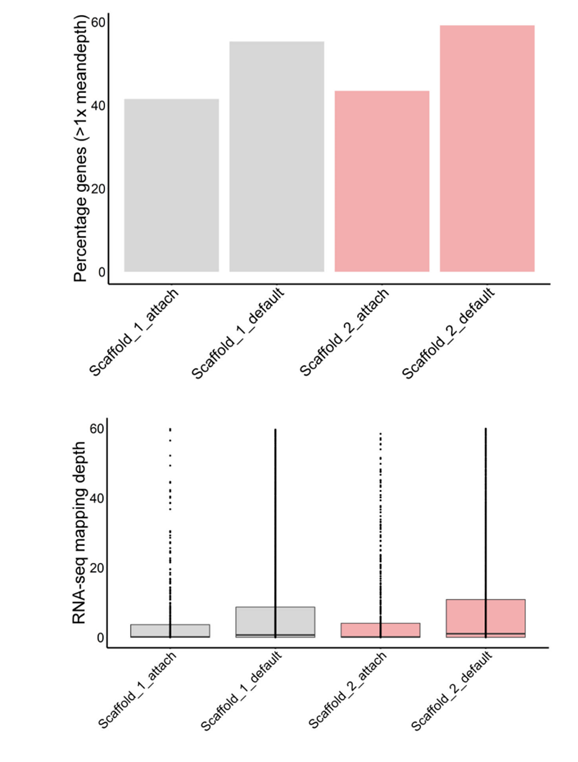

**Supplementary Figure 8**. The top panel shows the proportion of RNA-Seq data mapping to gene models in the manually incorporated versus remaining regions of super-scaffolds 1 and 2. The bottom panel shows the equivalent RNA-Seq mapping depth in these regions (cut-off at 60x mapping depth). The RNA-Seq data shown combines reads from heart, striated muscle and gonad. “Scaffold_1_attach” and “Scaffold_2_attach” on the x-axis represent the manually incorporated regions, while “Scaffold_1_default” and “Scaffold_2_default” represent the rest of super-scaffolds 1 and 2, respectively.
